# Switching on Behavioral and Neural Rhythmicity to Retrieve Memories When the Number of Retained Items Exceeds Four

**DOI:** 10.1101/2024.08.31.610596

**Authors:** Takuya Ideriha, Junichi Ushiyama

## Abstract

Even when we experience difficulty in recalling memories, we nevertheless manage to retrieve the target items. However, the neural mechanisms that enable such difficult memory retrieval are unknown. Here, we report an intriguing phenomenon where our nervous system “switches on” behavioral/neural rhythmicity to retrieve memory when the number of candidate items exceeds four. In our experiments, participants learned and retrieved 2–5 color/letter pairs. Analyses of hundreds of reaction times revealed a significant tendency for memory recall to occur at discrete times corresponding to theta–alpha (4–13 Hz) cycles, but only when the number of memorized pairs exceeded four. Electrophysiological data localized theta–alpha rhythmicity around parietal electrodes, a region associated with the long-term memory system. Our findings suggest that neural rhythmicity facilitates memory retrieval when the number of candidate items exceeds four, which is known as the “magical number” corresponding to the limit of human cognitive capacity.

## INTRODUCTION

We often experience difficulty recalling information that we have learned. This is especially true when studying a large amount of material, such as for a vocabulary test. Despite this difficulty, we nevertheless manage to retrieve these memories. How does our nervous system deal with a challenging memory recall scenario? Although there have been no direct investigations into this issue to our knowledge, some insights are provided from neurobiological research.

Previous studies have shown that rhythmic neural activity, called neural oscillations, has essential roles in memory recall [1,2]. Of great interest are theta–alpha (4–13 Hz) oscillations around the brain regions comprising the default mode network, including the hippocampus, medial prefrontal cortex, posterior cingulate cortex, and angular gyrus [3–5]. For example, a human high-density electroencephalography (EEG) study showed that the neural activation pattern of items to be recalled emerged specifically in the 8 Hz rhythm phase in the hippocampus [6]. Additionally, theta– alpha oscillations are reported to be synchronized across brain regions during episodic memory retrieval [5,7,8]. Watrous et al. demonstrated that the synchronous low-frequency (1–10 Hz) activity seen among default mode network regions was stronger during successful memory recall compared with incorrect trials [9]. These reports suggest the importance of neural rhythmicity in memory recall, emphasizing the need for deeper investigations of when and how rhythms contribute to memory retrieval.

To investigate the relationship between neural rhythmicity and cognitive functions, analysis of “behavioral rhythmicity” is among the most promising methods [10–13]. Of great relevance to our study, ter Wal et al. performed an associative memory task involving the memorization and recall word/image pairs [14]. The distribution of reaction times (RTs) for recall items over hundreds of trials showed multimodality, corresponding to a rhythmicity of about 4 Hz (an approximately 250-ms interval between peaks), consistent with the theta rhythm in neurophysiology. This result suggests that the theta neural oscillations seen during memory recall are robust enough to be observed at the behavioral level. Behavioral rhythmicity is a surprising phenomenon that was revealed purely through RT analysis. These behavioral methods possess advantages that complement neurophysiological methods [15]. Because neurophysiological data contain a vast amount of information on neural activity and noises that are not critically important to the task at hand, it is challenging to determine which neural activities are meaningful. In contrast, behavioral outputs reflect the final outcome of neural processing, such that the obtained data are inevitably of critical relevance to task execution. Therefore, behavioral and neurophysiological data have historically complemented each other, advancing our understanding of neuroscience by compensating for each other’s weaknesses.

However, it remains unclear how our nervous system handles challenging memory recall situations, such as when many items are retained in memory and/or when they are not well consolidated. Does the brain use a unified mechanism to recall memories regardless of whether the recall is easy or difficult? Here, by leveraging both behavioral and neurophysiological methodologies, we report an intriguing phenomenon whereby our nervous system “switches on” rhythmicity when the number of candidate items exceeds four, and as the memories are consolidated. The number four has long been known to represent a cognitive boundary, i.e., it corresponds to the limit of human short-term memory and attention; as such, it is known as the “magical number four” [16]. Our results suggest that this invisible boundary is also applicable to non-short-term memory retrieval, and rhythms are used to overcome this limitation.

## MATERIALS AND METHODS

### Ethical approval

All experiments were conducted in accordance with the Declaration of Helsinki, except that the study was not pre-registered in a database and approved by the Research Ethics Committee at Shonan Fujisawa Campus, Keio University (receipt number 293). The participants received sufficient explanations about the experimental purpose and methods and provided written informed consent before participation.

### Participant details

We conducted two experiments. In Experiment 1, 15 participants (mean ± standard deviation [SD] age, 21.1 ± 2.0 years; 8 males, 7 females [1 left-handed]) completed 2–4-pair conditions (see Experimental design section for details), and a further 15 participants (mean ± SD age, 23.8 ± 2.7 years; 8 males, 7 females [1 left-handed]) completed a 5-pair condition. In Experiment 2, 25 participants (mean ± SD age, 25.2 ± 3.7 years; 17 males, 8 females [2 left-handed]) completed both the 3- and 5-pair conditions. All participants had normal or corrected-to-normal visual acuity and reported no color vision deficiency.

### Experimental design

Throughout all the experiments, the stimuli were generated, and the program run, using MATLAB (MathWorks, Natick, MA, USA) and the Psychophysics Toolbox Version 3 [17]. The monitor was positioned approximately 57 cm from the participant. Experiments 1 and 2 were identical, except that EEG was recorded simultaneously in Experiment 2. There were 2–5-pair conditions in Experiment 1, whereas only 3- and 5-pair conditions were included in Experiment 2.

In the experiment, participants performed a task wherein they learned color/letter pairs (Fig. 1a). In each trial, a colored circle was presented in the center of the screen for 1,500 ms, followed by the corresponding letter. The number of pairs to remember in each set varied by condition, ranging from two to five pairs. Each pair appeared 20 times per set, allowing participants to gradually form a strong memory. The order of stimulus presentation was randomized. Participants were instructed to type the corresponding letter on the keyboard accurately and quickly when a color appeared. If they could not recall the letter pair (including the first encounter with the color), they were instructed to wait for 1,500 ms and then rememorize the corresponding letter. Conversely, if they remembered the letter pair, they were asked to respond as quickly as possible within the 1,500 ms during which the colored circle was displayed. When participants responded within 1,500 ms, feedback indicating whether they were correct or incorrect quickly appeared on the screen. In instances of correct responses, the corresponding letter was displayed in blue, whereas for incorrect responses, the correct letter was displayed in red for 500 ms, serving as feedback. If a response was not provided within 1,500 ms, the correct letter was displayed, and participants were instructed to memorize the color/letter pair again and press the key. After a 1,000 ms inter-trial interval, the next trial began. A specific color/letter pair appeared 20 times within a single set. Thus, in the 3-pair condition, for example, there were 60 (3 × 20) trials per set, and the order of presentation of the pairs was randomized. Participants learned color/letter pairs through such sets and gradually responded more quickly. The number of trials and sets per condition are listed in Table 1. Although the color/letter pairs were fixed within each set, they changed between sets. Consequently, participants were required to learn new pairs for each set. Participants could take breaks between sets. The color codes for the eight candidate colors were in RGB format: [230,0,18], [243,152,0], [207,219,0], [34,172,56], [0,160,233], [29,32,136], [146,7,131], and [228,0,127]. The candidate letters for the key inputs were “s”, “d”, “f”, “j”, “k”, and “l”. In each set, colors and letters were chosen at random from among the candidates. The fingers used to make responses were the left and right ring fingers, middle fingers, and index fingers.

**Table 1.**
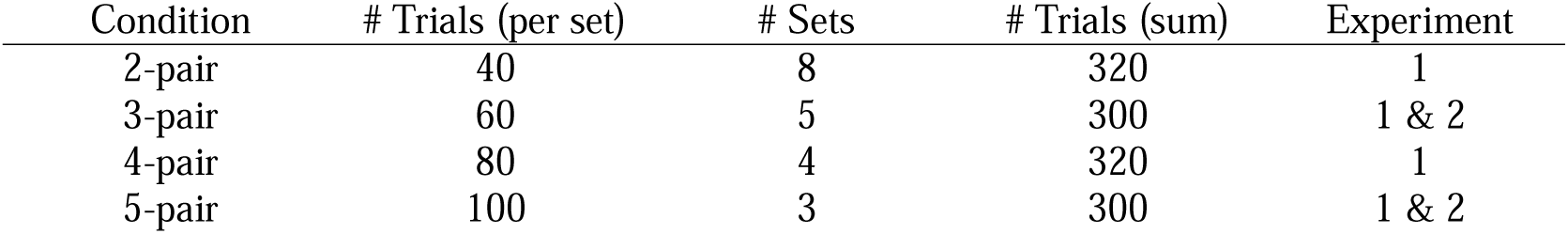
Experimental conditions and trial counts.

| Condition | # Trials (per set) | # Sets | # Trials (sum) | Experiment |
| --- | --- | --- | --- | --- |
| 2-pair | 40 | 8 | 320 | 1 |
| 3-pair | 60 | 5 | 300 | 1 & 2 |
| 4-pair | 80 | 4 | 320 | 1 |
| 5-pair | 100 | 3 | 300 | 1 & 2 |

**Figure 1.**
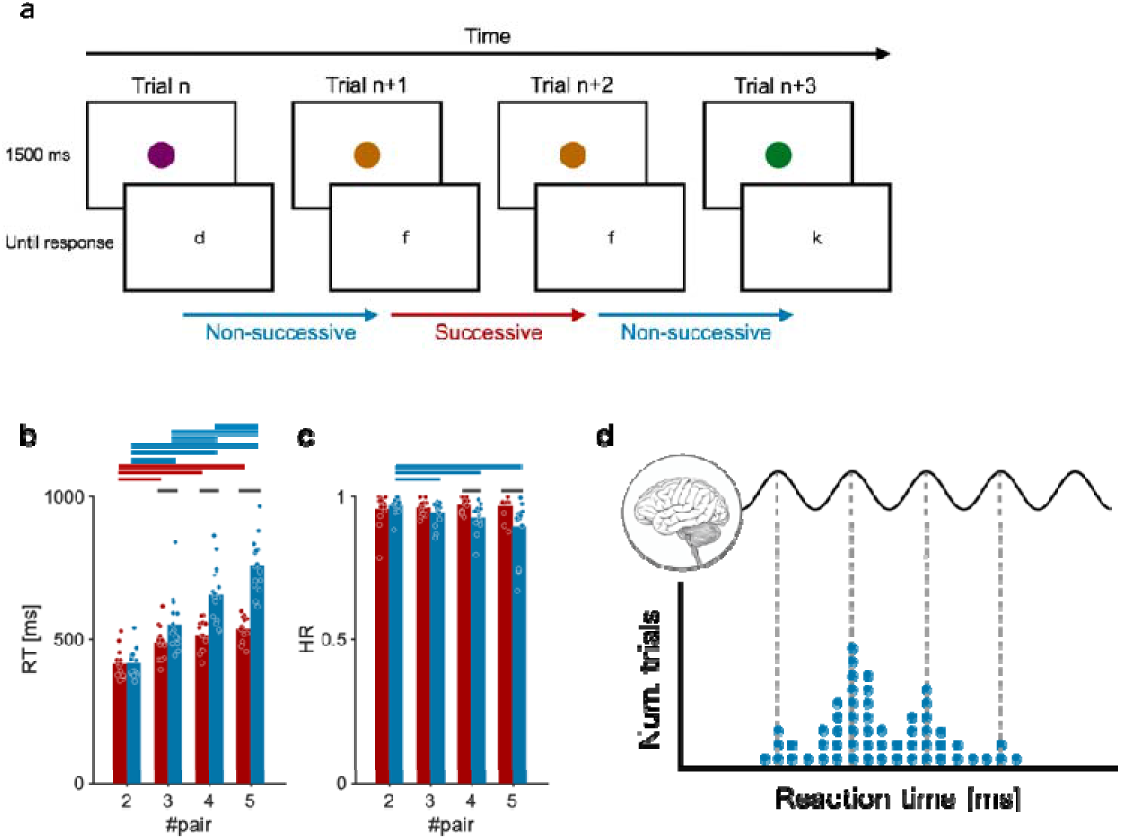
(a) Task procedure. According to the memorized color/letter pairs, participants quickly pressed the corresponding key upon the appearance of a colored circle. Successive trials denote trials with the same color as the previous trials; all other trials are non-successive trials. (b) The average reaction time (RT) in all conditions. Each dot denotes the average RT of an individual participant. Data from successive trials are in red, whereas data from non-successive trials are in blue. The horizontal bars above denote significant differences (p < 0.05). (c) The average hit rate in all conditions. (d) Schematic illustration of the relationship between neural oscillation and behavioral rhythmicity.

### EEG recordings

Scalp EEG was recorded from 19 scalp positions on the basis of the International 10–20 system in Experiment 2. Ag/AgCl electrodes with a diameter of 18 mm (g.LADYbirdPASSIVE 1035; Guger Technologies, Graz, Austria) were affixed to an EEG cap (g.GAMMAcap 1027; Guger Technologies) covering the whole head. Reference and ground electrodes were placed on the left and right earlobes, respectively. EEG was amplified and band-passed (0.5–200 Hz) via an analog biosignal amplifier (g.BSamp 0201A; Guger Technologies). Analog EEG signals were converted into digital signals by an analog-to-digital converter (NI USB-6259; National Instruments, Austin, TX, USA), at a sampling rate of 1,000 Hz, controlled by a data-logger program designed using MATLAB software (MathWorks).

### Behavioral data analyses

#### Trials included

Only data from correct trials (correct responses within 1,500 ms) were analyzed. Additionally, we analyzed trial data after all pairs had appeared at least once. For example, if there were pairs A–C in the 3-pair condition and the order of trials was “A, B, B, A, C, B…”, trials after the sixth trial (B) were included in the analysis. For each participant, the analysis of each condition was performed by merging the data across all sets within the same condition. For example, in the 3-pair condition with five sets of 60 trials, data from 300 trials were used in the same analysis. Of these, data that met the above criteria, i.e., correct responses within 1,500 ms after all pairs had appeared at least once, were analyzed according to the procedure described in the following. The average hit rate (HR) and RT were also calculated from these trials.

#### Successive and non-successive trials

In each condition, we divided trials into “successive” and “non-successive” trials. Successive trials are those having stimuli of the same color as in the previous trials, and all other trials are non-successive trials. Because the data from successive trials are easily recalled and likely to exhibit characteristics different from those of the other trials (Fig. 1b), we analyzed them separately.

#### RT distribution analyses

Of the RT data that met the criteria above, those exceeding three median absolute deviations were excluded from the analysis as outliers. RT distributions were calculated using the kernel density estimation method by sliding a 50-ms rectangular window ranging from 1 to 1,500 ms with a 1-ms resolution. The obtained distribution was smoothed by a moving average using a 20-ms window. To analyze the time-resolved frequency representation in the non-stationary time course of the RT distributions, we applied Morlet wavelet decomposition using the “cwt” function of the Wavelet Toolbox for MATLAB. The above method was used to assess the oscillatory characteristics of the behavioral rhythmicity instead of the Fourier transform, because the wavelet decomposition does not assume stationarity of the time courses. The average value in the 4–13 Hz range over the entire time course of the obtained decomposition was calculated as the quantitative theta–alpha value. This value was Z-scored together with the results of the surrogate analyses described below and used for statistical analysis. We defined this value as the “behavioral theta–alpha value”.

#### Surrogate analyses

To examine the existence of theta–alpha oscillations in the RT distributions, we generated null-hypothesis surrogate distributions for each participant as follows. First, we fitted an ex-Gaussian distribution to the original RT distribution. Generally, an ex-Gaussian distribution fits a human RT distribution well and has been used to examine human response characteristics, including the retrieval of memories [18–20]. Then, a number of simulated RTs equal to the number of corresponding trials were randomly sampled from the ex-Gaussian distribution. After that, the same frequency analyses as performed originally were conducted for the simulated RTs. This process was repeated 500 times, and the mean and SD of the null-hypothesis data were obtained. Using these data, the original theta–alpha value was translated into a Z-score.

#### Comparison between the early and late trials

To test the effect of learning on rhythmicity, we compared the data of non-successive trials from the 5-pair condition between the early and late trials in each set. We first extracted non-successive trials on the basis of several criteria (see the Trials included section). The first and last 10 trials of each set were extracted, resulting in a total of 30 first and 30 last trials (because each participant completed three sets of the 5-pair condition). Note that the above criteria reflected how well each memory was consolidated because participants should have learned new pairs in each set. Data from participants with insignificant behavioral theta–alpha values (peak value in the theta– alpha range < 1.96; p > 0.05) were excluded from further analyses. To ensure a sufficient number of participants in this analysis, we aggregated data from Experiments 1 and 2, with 17 participants showing significant behavioral oscillations. For each participant, the peak frequency in the 4–13 Hz range of the oscillations was calculated, and a bandpass filter was applied in the 0.5 Hz range before and after the peak frequency, resulting in a 1 Hz range (second-order Butterworth filter). A Hilbert transform was applied to the obtained waveform to extract the phase of the oscillation. Phase 0 corresponds to the rhythmic peak of the behavioral oscillations. Using this, we calculated the phase where the response was made in a certain trial. The V-test was used to determine whether the RTs of the first 30 and the last 30 trials were biased toward phase 0 of the oscillation. These V-tests were performed for each individual. In addition, phase data for the first and the last 30 trials were aggregated across individuals, and whether the phase was biased toward 0 degrees was also determined using V-tests. This analysis calculated how strictly the reactions in the first and last 30 trials extracted were phase-locked to the behavioral oscillations. After that, we statistically compared the first 30 and last 30 trials. However, the results from the above-mentioned analysis should be interpreted carefully because we could not avoid circularity: the data from trials used to calculate the phases of the reactions were also included in the calculation of RT distributions. Therefore, we conducted an additional analysis excluding the first 30 and last 30 trials extracted from the calculation of the RT distribution. In this additional analysis, the number of participants who showed significant behavioral oscillations decreased from 17 to 14 because fewer trials were used for the calculation of RT distributions. Hence, data from these 14 participants were included in the statistical analysis. Other processes were carried out in the same manner as in the aforementioned analysis.

#### Comparison between correct and the incorrect trials

We examined whether the RTs in trials with incorrect responses made within 1,500 ms were phase-locked to the behavioral oscillations of correct trials. As inclusion criteria, data from participants with significant oscillations in correct trials (peak Z-score for behavioral rhythmicity in the theta–alpha range > 1.96; p < 0.05) and ≥ 10 trials with incorrect responses made within 1,500 ms were chosen for further analyses. Trials with incorrect responses made within 1,500 ms are those in which the participants recalled and answered with incorrect letters, rather than simply failing to recall them. After applying this inclusion criterion, only five participants remained in Experiment 1. We therefore decided to analyze the data of 13 participants by combining the data from Experiments 1 (n = 5) and 2 (n = 8). In the analysis, we first randomly extracted equal numbers of correct and incorrect trials. The RTs of correct trials that remained unextracted were used to calculate behavioral oscillations, to examine whether the RTs of the incorrect and extracted correct trials were locked to the phase of those oscillations and to avoid circularity. Similarly to the comparison between the early and late trials, the waveform of the peak frequency in the 4–13 Hz range was extracted with a bandpass filter (second-order Butterworth filter) in the 0.5 Hz range before and after the peak frequency. A Hilbert transform was applied to obtain the phase of the oscillation. Using this, we identified the phase where the response was made in a certain trial. The V-test was applied to determine whether the RTs of incorrect and extracted correct responses were biased toward phase 0 (peak phase) of the oscillation. These V-tests were performed for each individual. In addition, phase data for incorrect and extracted correct responses were aggregated across individuals, and whether the phase was biased toward 0 degrees was determined using V-tests.

### EEG data analyses

#### EEG preprocessing

Recorded EEG signals were processed with MATLAB and the EEGLAB toolbox (version 2023.0) [21]. The EEG signals were high-pass filtered at 3 Hz, and band-stop filtered at 49–51, 99– 101, and 149–151 Hz, using a third-order Butterworth filter. Data segments with visible noise caused by body movement, muscle activity, or poor contact were visually identified and excluded from the analysis. The remaining data were decomposed through independent component analysis, and components with eye blinks and eye movements were removed using the “runica” function of EEGLAB [21]. The data were then separated into epochs to calculate event-related potentials (ERPs). For stimulus-locked potentials, the epoch length was 6 seconds, spanning the period from 2 seconds before to 4 seconds after the stimulus. For reaction-locked potentials, 6-second epochs were generated, covering the 4 seconds before and 2 seconds after the reaction.

#### EEG analyses

In this study, we calculated ERPs to examine the stimulus/reaction-locked rhythmicity of EEG. In the EEG analyses, we only analyzed correctly answered non-successive trials because the number of successive or incorrect trials would not have been sufficient for calculating ERPs. The ERP calculations were conducted within 500-ms time windows: from 0 to 500 ms post-stimulus for stimulus-locked potentials, and from −500 to 0 ms pre-reaction for reaction-locked potentials. Considering that both time windows were 500 ms, trials where the RT for a given stimulus was within 500 ms were excluded from the analysis. To unify the analysis conditions for the 3- and 5-pair conditions, the number of trials analyzed was aligned with the condition having fewer trials. For conditions with more trials, data were randomly sampled to match the number of trials in the condition with fewer trials. For instance, if a participant had 176 analyzable trials in the 3-pair condition and 120 in the 5-pair condition, the analysis was standardized to the 120 trials of the 5-pair condition. From the 3-pair condition, 120 trials were randomly selected for analysis, and the rest were excluded. The final number (mean ± SD) of epochs used in the analysis of each participant’s data was 112.3 ± 28.5 trials. For the calculation of ERPs, the data of each epoch were low-pass filtered at 50 Hz using a third-order Butterworth filter. The ERP analysis was performed for each individual electrode. Stimulus-locked potentials were calculated by summing and averaging the waveforms of each epoch aligned with the moment of stimulus presentation. Reaction-locked potentials were calculated by summing and averaging the waveforms of each epoch aligned with the moment of reaction. The stimulus/reaction-locked potentials resulted in 19 waveforms per participant because we recorded data from 19 channels. For each resulting ERP epoch, Morlet wavelet transform was applied using MATLAB’s cwt function, similar to the calculation of behavioral oscillations, and averaged over time for the target 500-ms period (from 0 to 500 ms for stimulus-locked potentials, and from −500 to 0 ms for reaction-locked potentials). Subsequently, values in the 4–13 Hz range were calculated, and the proportion of the theta/alpha band was obtained by dividing by the total value below 50 Hz. These values will be used as quantitative EEG metrics in the future. Furthermore, the peak frequency was defined as the frequency with the highest power within the 4–13 Hz range. We defined this value as the “EEG theta–alpha value”.

#### Statistical analyses

Unless stated otherwise, we applied non-parametric statistical measures because the data did not all meet the assumption of a normal distribution. When comparing data between two conditions, we adopted Wilcoxon’s signed-rank test. To examine the correlation between two variables, we adopted Spearman’s rank correlation test. Before applying two-way analysis of variance (ANOVA), we performed the aligned rank transform procedure [22], which enables us to apply an ANOVA to non-parametric data. When the interaction was significant, we applied Kruskal–Wallis tests to examine the simple main effects. After that, to conduct a statistical comparison among several conditions, we applied the Wilcoxon rank-sum test, correcting the p-value with Shaffer’s method [23], and to conduct a statistical comparison between two conditions, we applied Wilcoxon’s signed-rank test. When comparing the theta–alpha power in EEG between the 3- and 5-pair conditions, we repeated the comparison for 19 electrode locations. Therefore, we controlled the false discovery rate with the Benjamini–Hochberg correction [24] by setting the q-value at 0.05. When examining whether the phase at the moment that a response was made was biased toward the 0-degree phase of the behavioral oscillations, a V-test was applied using the CircStat toolbox [25] in MATLAB.

## RESULTS

### Overall task performance

In the experiment, participants were required to remember color/letter pairs and answered with the corresponding letter quickly when a colored circle appeared on the screen (Fig. 1a). Essentially, task performance was worst when participants were required to recall non-successive items from among a large amount of information (Fig. 1b, c). The two-way ANOVA of RT showed significant main effects of condition (the number of retained pairs) (F[3, 112] = 69.74, p < 0.001) and successiveness (whether the trial was successive or non-successive) (F[1, 112] = 86.05, p < 0.001), and the interaction was also significant (F[3, 112[ = 17.24, p < 0.001). Thereafter, we conducted Kruskal–Wallis tests to examine the simple main effect of condition. In both successive and non-successive trials, the results showed a significant simple main effect of condition (successive: p < 0.001; non-successive: p < 0.001). Regarding successive trials, Wilcoxon rank-sum tests (Shaffer-corrected) showed that the RT in the 2-pair condition was shorter than those in the 3– 5-pair conditions (p < 0.001), and the RT in the 3-pair condition was shorter than that in the 5-pair condition (p = 0.029). The other comparison showed no significant differences (p > 0.05). Regarding non-successive trials, the same tests showed that all comparisons were significant, and the RT in larger-pair conditions was longer than that in smaller-pair conditions (p < 0.001 in 2- and 3-pair, 2- and 4-pair, 2- and 5-pair, and 3- and 5-pair comparisons; p < 0.01 in 3- and 4-pair and 4- and 5-pair comparisons). Additionally, we performed four Wilcoxon signed-rank tests to examine the simple main effect of the successiveness of trials on RTs. The results showed no significant difference in the 2-pair condition (p > 0.05), but the RTs in non-successive trials were longer than those in successive trials in the 3–5-pair conditions (p = 0.0015 in the 3-pair condition; p < 0.001 in the 4- and 5-pair conditions). The two-way ANOVA of the HR showed significant main effects of condition (the number of retained pairs) (F[3, 112] = 3.47, p = 0.019) and successiveness (whether the trial was successive or non-successive) (F[1, 112] = 21.08, p < 0.001), and the interaction was also significant (F[3, 112] = 6.85, p < 0.001). Thereafter, we conducted Kruskal–Wallis tests to examine the simple main effect of condition. In successive trials, the simple main effect of condition was not significant (p > 0.05), whereas in non-successive trials, the simple main effect of condition was significant (p < 0.001). Regarding non-successive trials, Wilcoxon rank-sum tests showed that the HR in the 2-pair condition was higher than those in the 3-pair condition (p = 0.012), 4-pair condition (p = 0.0013), and 5-pair condition (p < 0.001), but no other comparison showed significant differences (p > 0.05). We also performed four Wilcoxon signed-rank tests to examine the simple main effect of the successiveness of trials on the HR. The results showed no significant difference in the 2- or 3-pair condition (p > 0.05), but the HR in non-successive trials was lower than those in successive trials in the 4- and 5-pair conditions (p = 0.0012 and p < 0.001, respectively). These results demonstrate that the retrieval from memory of non-recently encoded items from among a large amount of information was the most difficult type of retrieval.

### Memory items were retrieved rhythmically in the 5-pair condition

Next, we visualized individual RT distributions and found behavioral rhythmicity in the distributions in non-successive trials of the 5-pair condition. Four typical RT distributions for each memory load condition are shown in Fig. 2a. We found clear periodicity in the RT distributions, especially for the RTs to non-successive stimuli in the 5-pair condition. To examine the oscillatory characteristics of the RT distributions, we conducted a time–frequency decomposition and compared them with the non-oscillatory surrogate RT distributions (500 iterations). The obtained mean and SD were used to calculate the Z-score of the oscillatory value. Fig. 2b and c show a clear peak around the theta–alpha band in the non-successive trials of the 5-pair condition. The Z-score of theta–alpha value was significantly larger than zero in the non-successive trials of the 5-pair conditions (Fig. 2d; one-tailed Wilcoxon signed-rank test, p = 0.0034), although significant differences from zero were not seen in any other trials/conditions (p > 0.05). These results suggest that memory retrieval becomes rhythmical when the number of retained items exceeds four. Below, we describe the results of the analysis of the behavioral rhythmicity of the non-successive trials of the 5-pair condition.

**Figure 2.**
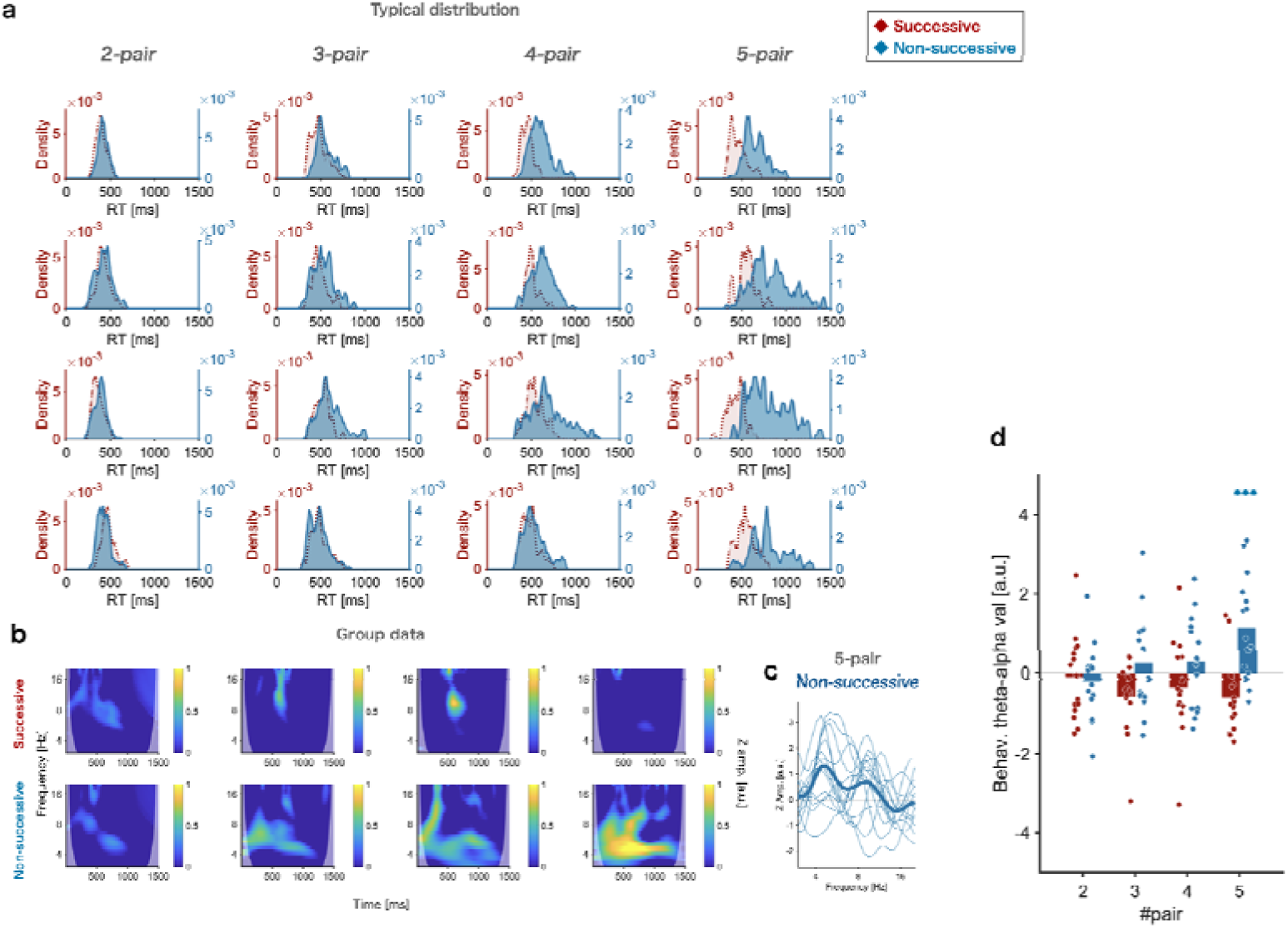
(a) Four typical RT distributions for each condition. Note that data in the same row are not from the same participant. The number of pairs increases from the left to the right column. The red color indicates data from successive trials, whereas the blue color indicates data from non–successive trials. (b) Time–frequency decomposition of the RT distribution averaged across participants. Each voxel indicates the behavioral theta–alpha value (Z–score of behavioral rhythmicity in the theta–alpha range). Figures in the upper row are from successive trials, whereas those in the lower row are from non–successive trials. (c) Frequency power of the RT distribution of non–successive trials in the 5–pair condition. Each line represents a single individual. (d) Behavioral theta–alpha value averaged across participants. Each dot denotes a single participant.

### Examining the relationship between behavioral rhythmicity and memory performance

We examined the inter-individual relationship between theta–alpha rhythmicity and task performance in non-successive trials of the 5-pair condition, where significant theta–alpha rhythmicity was observed. Interestingly, the theta–alpha values were not correlated with either the HR (rho = 0.039, p = 0.89) or the average RT (rho = −0.33, p = 0.23). Additionally we examined the correlation between the peak frequency of behavioral rhythmicity and task performance, observing no significant correlation (HR: rho = 0.29, p = 0.30; RT: rho = −0.067, p = 0.81). These results suggest that the subjective difficulty of memory recall and prolonged RTs themselves are not determinants of rhythmicity in memory recall at the inter-individual level. In addition, although peak frequency showed individual differences in the theta–alpha band, this would not be related to memory performance.

### Later trials showed stronger phase locking to behavioral rhythmicity

Does this rhythm appear when the number of items exceeds 4 or when recall is difficult because of insufficient learning? Here, we combined data from participants who showed significant behavioral oscillations in Experiments 1 and 2 to ensure that a sufficient number of participants were included, resulting in 17 participants’ data being analyzed. In a typical RT distribution, RTs from 30 late trials seem to be more strongly clustered around the peaks of the RT distribution than those from 30 early trials (Fig. 3a). When the average RT was calculated for each 30-trial bin, the RTs showed a decreasing trend as the trials progressed, indicating that a learning process was occurring in the participants (Fig. 3b). In contrast, the V-value, which reflects the extent to which reactions were biased toward phase zero (peak phase) of behavioral rhythmicity, showed an increasing trend as trials progressed, indicating that the reactions were locked to the behavioral rhythmicity to a greater degree as participants learned the memory pairs. To statistically examine this trend, we compared the averaged RT and rhythmicity between the first 30 and last 30 trials. The average RT was significantly shorter in the last 30 trials among non-successive trials of the 5-pair condition (Fig. 3b, Wilcoxon’s signed-rank test, p = 0.0017), showing that participants could retrieve memory items more easily in the last 30 trials compared with the first 30 trials. Importantly, reactions in the last 30 trials showed significantly larger V-values than those in the first 30 trials (Fig. 3a, c, Wilcoxon’s signed-rank test p = 0.0014). This indicates that reactions in the late trials showed a stronger association with rhythmic recall than those in earlier trials. The V-values for both the first (Wilcoxon’s signed-rank test p = 0.0031) and the last (Wilcoxon’s signed-rank test p < 0.001) 30 trials were significantly larger than 0. When aggregating the phases of 30 trials across the 17 individuals, which resulted in 510 reactions each for the first and the last trials, the phases of both the first 30 (V = 45.67, p = 0.0021) and the last 30 (V = 128.24, p < 0.001) trials were significantly biased toward 0 degrees (Fig. 3f).

**Figure 3.**
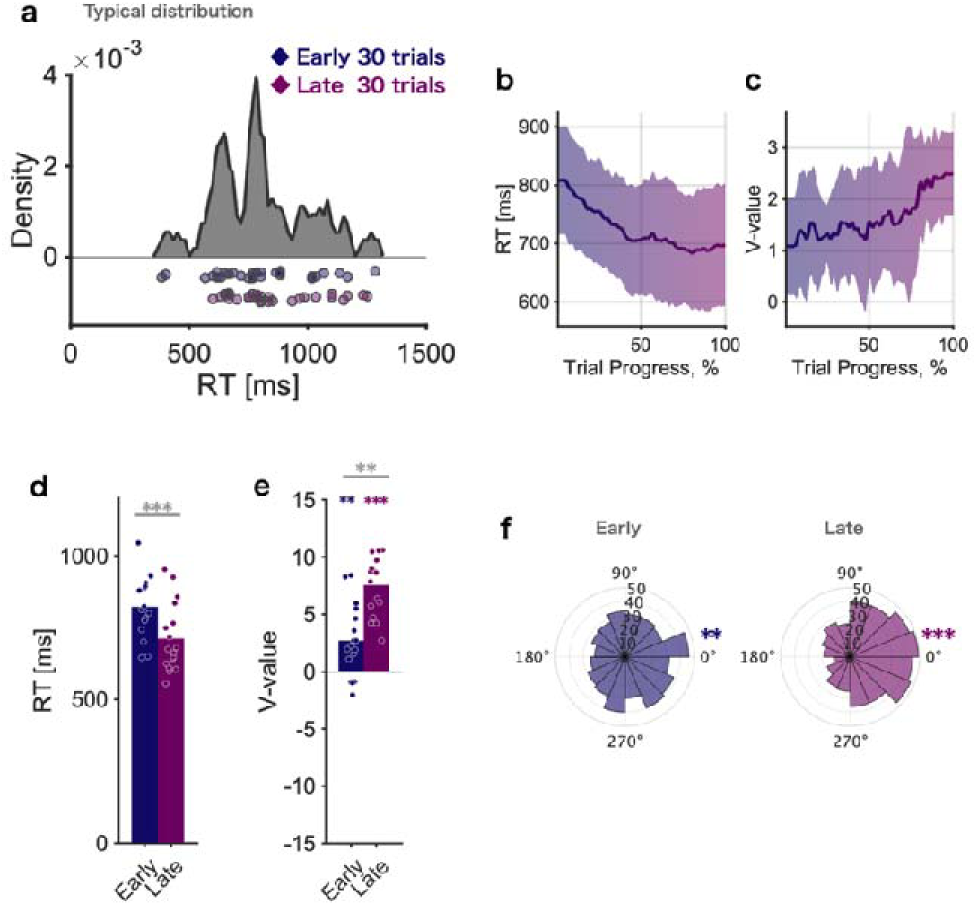
(a) RT distribution for a typical participant. Scattered dots denote RTs from 30 early trials (upper row) and 30 late 30 (bottom row). (b) Change in RT according to trial progress averaged across individuals. The shaded area denotes the standard deviation (SD). (c) Change in Vvalue according to trial progress averaged across individuals. The shaded area denotes the SD. (d) Averaged RTs of 30 early and 30 late trials. Each dot denotes a single participant. (e) Averaged Vvalue of 30 early and 30 late trials. Each dot denotes a single participant. (f) Circular histogram of trials aggregated across participants denoting the phase where each response was made. The left panel is for 30 early trials, whereas the right panel is for 30 late trials.

In these analyses, we examined the contributions of the first 30 and last 30 trials to the rhythmicity of the RT distribution. For this purpose, the 60 extracted trials were also included when creating the RT distribution. However, this approach may overestimate the degree to which the extracted trials are locked to the phase of the RT distribution’s rhythm because of the influence of circularity. To address this concern, we also conducted analyses excluding the first 30 and last 30 extracted trials from the RT distribution. As shown in Fig. 3e, whereas the V-value for the last 30 trials was again significantly larger than 0 (Wilcoxon’s signed-rank test p = 0.0040), the V-value for the first 30 trials was not (Wilcoxon’s signed-rank test p = 0.90). The difference in V-values between the first 30 and last 30 trials was not significant (Wilcoxon’s signed-rank test p = 0.058). In aggregated phase analysis, whereas the V-value for the last 30 trials was again significantly larger than 0 (V = 42.54, p = 0.0017), the V-value for the first 30 trials was not (Fig. 3f, V = −1.52, p = 0.54) (Fig. 3f). These results suggest that this rhythmicity does not emerge when memory recall is difficult because of insufficient learning but does emerge if retrieving a memory when at least five items have been well consolidated.

### Incorrect trials were not phase-locked to behavioral rhythmicity

Next, we examined whether or not trials with incorrect responses made within 1,500 ms were phase-locked to the oscillations of correct trials among the non-successive trials of the 5-pair condition. We chose data from participants with significant behavioral oscillations and ≥ 10 trials with incorrect responses made within 1,500 ms among the non-successive trials of the 5-pair condition. Because these criteria led to only 5 participants remaining in Experiment 1, we combined the data from Experiments 1 and 2, yielding 13 participants. By avoiding circularity (see Materials and Methods), we could analyze whether correct and incorrect trials were biased towards the peak phase of the recall rhythm. As shown in Fig. 4a, the V-value (non-uniformity around 0°) was significantly larger than 0 in correct trials (Wilcoxon’s signed-rank test, p = 0.017), but not in incorrect trials (p = 0.45). As an aggregated phase analysis (Fig. 4b, c), RT again tended to be biased toward 0 degrees (peak phase) in correct trials (V = 26.28, p = 0.012), but not in incorrect trials (V = 14.50, p = 0.11). Although the results should be interpreted carefully because of the relatively small samples, they suggest that incorrect responses are not locked to behavioral rhythmicity.

**Figure 4.**
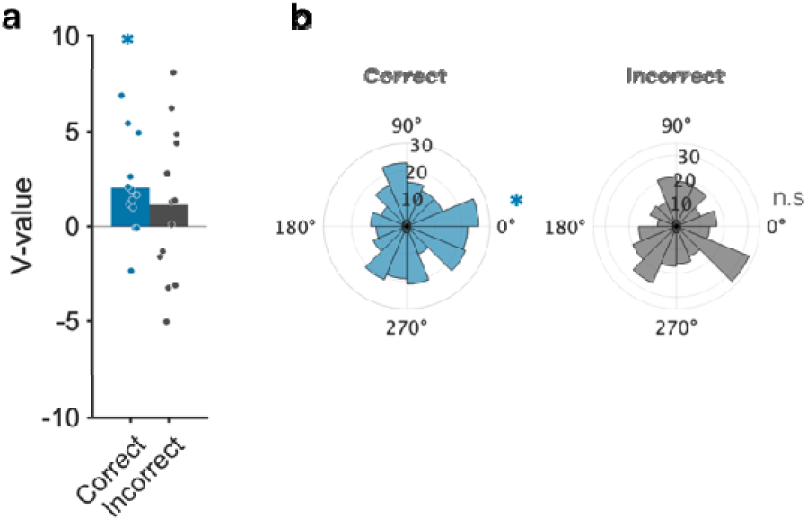
(a) Average V-value for correct and incorrect trials. Each dot denotes a single participant. (b) Circular histogram of trials aggregated across participants denoting the phase where each response was made. The left panel shows correct trials, and the right panel shows incorrect trials.

### Replication of behavioral rhythmicity in Experiment 2

In Experiment 2, 25 participants completed the 3- and 5-pair conditions while neural activity was recorded with EEG. First, we replicated the behavioral rhythmicity seen in Experiment 1 (Fig. 5). Again, we found that the theta–alpha value of behavioral rhythmicity in the non-successive trials of the 5-pair condition was significantly larger than 0 (Fig. 5a, b, one-tailed Wilcoxon signed-rank test, p = 0.0062), but not in the successive trials and/or the 3-pair condition (p > 0.05). The theta–alpha values were not correlated with either the HR (rho = 0.055, p = 0.79) or the average RT (rho = −0.34, p = 0.095) at the inter-individual level. Peak frequency did not correlate with either the HR (rho = 0.14, p = 0.50) or the average RT (rho = 0.13, p = 0.55) across individuals. These results are consistent with the behavioral results of Experiment 1, supporting the existence of behavioral theta–alpha rhythmicity in the non-successive trials of the 5-pair condition.

**Figure 5.**
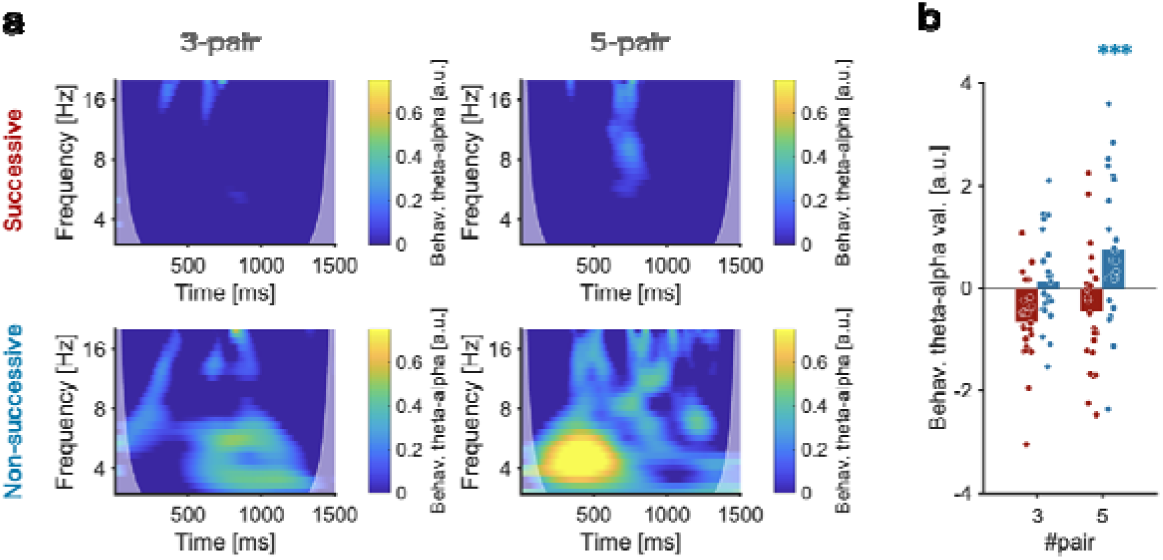
Replication of behavioral results. (a) Time–frequency decomposition of the RT distribution averaged across participants. Each voxel indicates a behavioral theta–alpha value. Figures in the upper row are from successive trials, whereas those in the lower row are from non–successive trials. (b) Behavioral theta–alpha value averaged across participants. Each dot denotes a single participant.

### Stimulus-locked potentials of EEG

The behavioral rhythmicity in the RT distribution implies the presence of phase-locked theta–alpha oscillations at the moment of stimulus presentation. Therefore, we calculated the theta– alpha power of the stimulus-locked potentials of EEG signals and compared them between the non-successive trials of the 3- and 5-pair conditions (Fig. 6a–c). Successive and incorrect trials were not analyzed because of the small number of trials available for EEG analyses. In a typical individual, the EEG trace showed more obvious theta–alpha rhythmicity in the 5-pair condition compared with the 3-pair condition around the parietal cortices (Fig. 6a, b). The difference in theta–alpha power between the 3- and the 5-pair conditions is shown in Fig. 6c. Theta–alpha power was stronger in the 5-pair condition than in the 3-pair condition around the lateral parietal electrodes (Fig. 6c). After adjusting the statistical threshold with the Benjamini–Hochberg method, the theta–alpha power was significantly stronger in the P7 (left lateral parietal) electrode (Wilcoxon’s signed-rank test, corrected p = 0.025). These results suggest that the rhythmic retrieval process in the 5-pair condition can be detected from the surface EEG signals and mostly depends on parietal regions.

**Figure 6.**
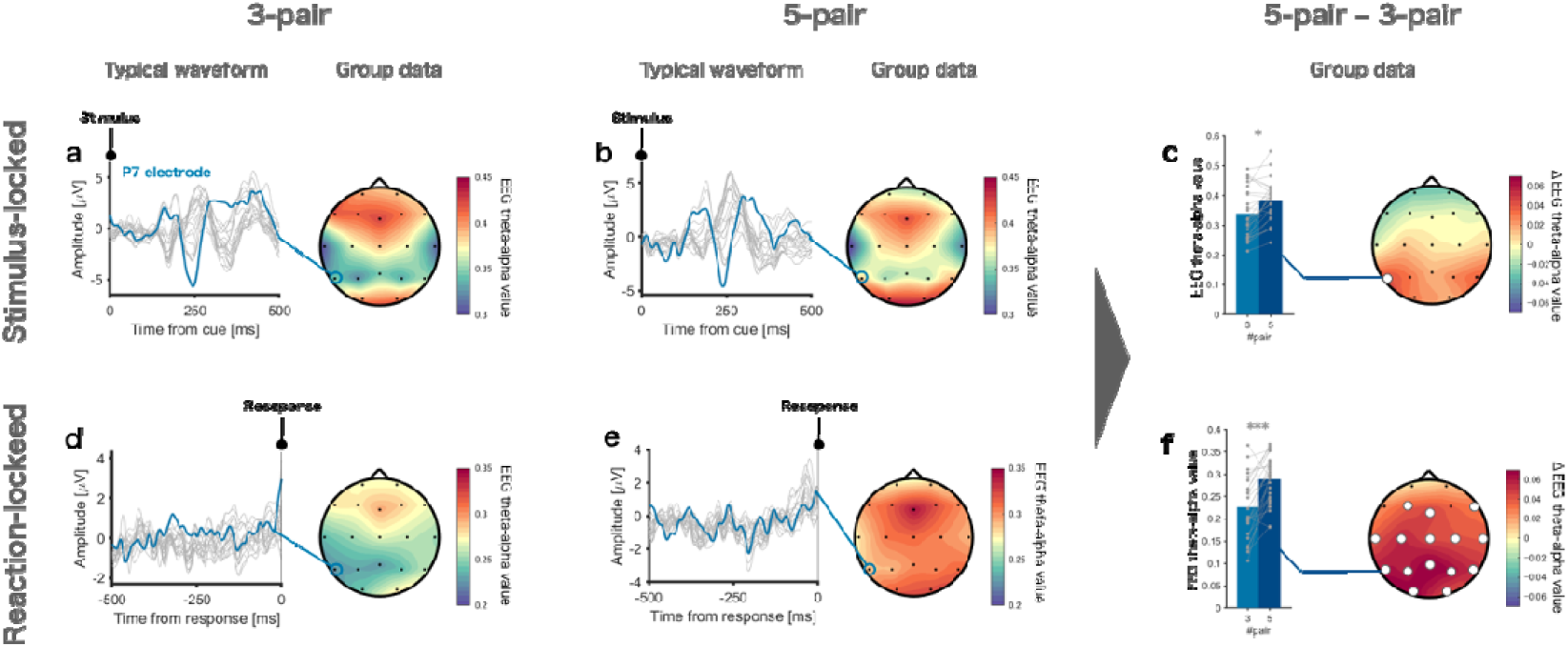
(a) Left: Waveforms of stimulus-locked potentials in the non-successive trials of the 3-pair condition from a typical participant. The blue line indicates a waveform of the P7 electrode. Each gray line indicates the waveform of a single electrode. Right: Grand average topography indicating the EEG theta–alpha value across all participants. (b) The same in the 5-pair condition. (c) Left: Bar graphs indicating the EEG theta–alpha value of the P7 electrode in the 3- and 5-pair conditions. Each dot and line denotes a single participant. Right: Grand average topography indicating the difference in EEG theta–alpha value between the 5- and 3-pair conditions. The white-filled circle indicates an electrode with a significant difference (Wilcoxon’s signed-rank test, corrected p < 0.05). (d–f) The same in the reaction-locked potentials.

### Reaction-locked potentials of EEG

Stimulus-locked potentials of EEG could be contaminated by several potentials, such as P300 [26] and P1 [27]. To circumvent any influence of these potentials and examine the rhythmicity in memory retrieval, we calculated the theta–alpha power in the reaction-locked potentials of EEG signals. This analysis examined the extent to which the reactions are locked to EEG rhythmicity. In a typical individual, we see clearer rhythmicity in the theta–alpha range in the 5-pair condition compared with the 3-pair condition (Fig. 6d, e). The difference in theta–alpha power between the 3- and 5-pair conditions is shown in Fig. 6f. Theta–alpha power was stronger in the 5-pair than in the 3-pair condition in most electrodes, as shown in Fig. 6f. Similar results between the stimulus- and reaction-locked potentials indicate the robustness of theta–alpha rhythmicity around parietal electrodes. The reaction-locked theta–alpha power showed wider topographic coverage than the stimulus-locked one, which may reflect the spread of the rhythm from the left lateral parietal lobe to the whole head over time.

## DISCUSSION

We investigated how the human nervous system achieves memory retrieval during difficult situations by combining behavioral and neural rhythmicity analyses. The novel findings are as follows: (1) in the 5-pair condition, the RT distribution showed distinct peaks, with the cycles corresponding to theta–alpha rhythmicity (Fig. 2); (2) in the 2–4-pair conditions, where recall performance was higher, the RT distribution showed no such significant rhythmicity; (3) even in the 5-pair condition, recall was not phase-locked to the behavioral rhythmicity in earlier trials where memory was not well consolidated, and the phase locking became stronger as the trials progressed and memory items were consolidated (Fig. 3); (4) in incorrect trials, recall was not significantly phase-locked to behavioral rhythmicity (Fig. 4); and (5) EEG analyses showed stronger theta–alpha stimulus/reaction-locked oscillations in the 5-pair condition than in the 3-pair condition, mostly around parietal electrodes (Fig. 6). Our results demonstrate that our nervous system switches on rhythmicity to retrieve consolidated memories when the number of retained items exceeds four.

Multiple peaks observed in the RT distributions for item recall suggest the involvement of neural oscillations in memory retrieval. Hundreds of memory recalls per participant in the 5-pair condition showed that memory recall tended to occur at specific/discrete times after memory cues, with the cycle corresponding to the theta–alpha range. As shown in Fig. 1, the discrete peaks in the RT distributions imply that the memory cue triggered phase reset of the neural oscillation [28], and the phase/frequency where phase resetting occurred was the same in almost all trials. Assuming that the phase of neural oscillation is the same across trials, we calculated stimulus-locked potentials of EEG signals across trials (Fig. 6): phase locking was mainly observed around the left parietal electrodes in the 5-pair condition, supporting the existence of phase resetting of neural oscillations triggered by the memory cue when the number of candidate items was five. In recent years, a growing body of research has investigated the nature of behavioral rhythmicity and gained insights into the neural mechanisms of attention [11–13,29], working memory [30–33], and long-term memory [14]. Adding to this body of research, our study is the first to demonstrate that behavioral rhythms associated with the retrieval of memories are switched on when the number of memorized items exceeds four.

We found an invisible boundary between numbers 4 and 5, which affects memory retrieval. As the term “ magical number 4” indicates, human cognitive ability has long been known to be severely limited when the number of items exceeds four [16,34]. In other words, the number of items we can actively hold in short-term memory at the same time [16,35], and the number of items we can pay attention to at the same time, is only about four [36]. Although fewer than five items can be instantly and accurately recognized, the identification of more than four items has been shown to rely on a vague estimation process [37,38]. It has been suggested that our nervous system possesses neurons that react specifically to 1–4 items, whereas there are no such neurons for higher numbers of items [38]. The current results are novel in showing that the non-short-term memory retrieval process is also affected by the cognitive boundary of four items. When the number of candidate items is within this boundary, retrieving them is an easy and smooth process, as reflected in the short RT, high HR, and lack of rhythmicity. Notably, when the number of candidate items exceeds four, such that the capacity for easy memory retrieval is limited, our nervous system apparently employs an alternative strategy, recruiting behavioral/neural rhythmicity. What is surprising about current results is that the non-short-term memory process was affected by the boundary, which has not been reported previously. Combined with previous reports, our results indicate that the human nervous system has a specialized neural infrastructure for processing four or fewer items.

Our results suggest that our nervous system recruits theta–alpha rhythmicity when the number of candidate memory items exceeds the working memory capacity of four, and items should be retrieved from long-term memory. Working memory is the short-term memory system that actively maintains and uses items that are “in mind” [39]. When the number of items is four or fewer, i.e., the above-described capacity of working memory, all candidate items can be actively retained in working memory storage throughout a memory task [16]. On the contrary, when the number of candidate items exceeds four, overflow of working memory capacity occurs, and the items should be stored in the long-term memory system to allow for memory retention and retrieval. The clear behavioral/neural rhythmicity seen in our 5-pair condition, where the items exceeded the working memory capacity, suggests the involvement of theta–alpha rhythmicity in long-term memory retrieval. Supporting this view, theta–alpha behavioral rhythmicity became progressively stronger in the latter trials, where memory items were well consolidated in long-term memory, whereas rhythmicity was not observed in earlier trials (Fig. 3). This indicates that the human nervous system possesses a highly flexible switching mechanism to overcome the limits of our cognitive capacity.

A recent study showed similar theta rhythmicity in a long-term associative memory task [14]. In that experiment, participants encoded more than 100 cue–target memory pairs and, after a distractor task (judging whether numbers displayed on a screen are even or odd), they retrieved the encoded items upon seeing a memory cue. In the task, which clearly depended on the long-term memory system, RT distributions showed behavioral rhythmicity, similar to our observations. On the basis of this result, the behavioral/neural rhythmicity observed in our tasks likely reflects recruitment of the long-term memory system. Notably, compared with the previous study, ours has two additional findings. First, our study found that the appearance of behavioral rhythmicity as a means to retrieve memories is situation-dependent: it appeared when the number of retained items exceeded four and when the items were well consolidated. Second, the frequency range was different between our study (4–13 Hz) and the previous one (1–5 Hz). This difference might indicate that the frequency reflects the difficulty of long-term memory retrieval: the more difficult the retrieval, the lower the frequency.

The EEG topography of theta–alpha oscillations in this study also supports the involvement of the long-term memory system in the 5-pair condition. The difference in theta–alpha oscillations between the 5- and 3-pair conditions was greatest around the left parietal electrodes (Fig. 6). These regions include posterior parietal cortices such as the angular gyrus, which is activated during episodic memory retrieval [3,40,41] and has connectivity with the hippocampus [9,42]. A possible function of theta–alpha neural oscillations is to synchronize neural activity across brain regions [43,44] related to episodic memory retrieval, namely the posterior parietal cortex, hippocampus, and other areas that are difficult to capture with surface EEG recordings. Using human intracranial recordings, Watrous et al. reported synchronization of the phase of slow oscillations (1–10 Hz) among the above-mentioned regions during long-term memory retrieval tasks. Therefore, the theta–alpha oscillations around parietal electrodes would indicate the synchronization of brain regions related to the long-term memory retrieval system, including the posterior parietal cortex and hippocampus.

The difference in EEG topography that we observed between stimulus- and reaction-locked potentials implies the propagation of theta–alpha oscillations from the left parietal cortex to other brain regions. While theta–alpha oscillations were observed around lateral parietal electrodes in the context of stimulus-locked potentials, they were observed around most electrodes, including in medial parietal, central, and frontal regions, in the context of reaction-locked potentials. Because reaction-locked potentials reflect later signals than those reflected by stimulus-locked potentials, the topographical differences in turn reflect the propagation of theta–alpha oscillations over time. In light of these considerations, long-term memory retrieval might be realized by synchronizing a wide range of brain regions, as triggered by theta–alpha oscillations in the left parietal region

In the current study, we observed behavioral/neural rhythmicity in the theta–alpha range when the number of candidate items to be retrieved from memory exceeded four. This rhythmicity likely reflects neural activity synchronized across brain regions related to long-term memory retrieval, a key mechanism for retrieving items when working memory capacity is exceeded. Our results might provide a useful basis to assess whether an item is well consolidated in long-term memory or is being dealt with in short-term memory. On the basis of our results, the status of stored memories can be further examined: an item retrieved through rhythmicity would be well consolidated in long-term memory, whereas an item retrieved without rhythmicity would not be well consolidated. Further work is needed to refine these findings and methodologies.

## ACKNOWLEDGEMENTS

This work was supported by grants from the Grant-in-Aid for Scientific Research (B) (Japan Society for the Promotion of Science, JSPS) (grant number 24K02845) to JU, a designated donation from Living Platform, Ltd, Japan to JU, and JST SPRING, Grant Number JPMJSP2123 to T.I. We thank Ms. Tomomi Hamaoka for their secretarial assistance and all other members of our laboratory for their insightful comments on the work. A portion of the sample in this study was recruited via a website to recruit experimental participants: https://www.jikkenbaito.com. We thank Michael Irvine, PhD, from Edanz (https://jp.edanz.com/ac) for editing a draft of this manuscript.

## AUTHOR CONTRIBUTIONS

Conceptualization, T.I. and J.U.; Methodology, T.I.; Sofware Development, T.I.; Data acquisition, T.I.; Data Analyses, T.I.; Visualization, T.I.; Interpretation, T.I. and J.U.; Writing—Original Draf, T.I.; Writing—Review & Editing, T.I. and J.U.; Supervision, J.U.; Funding Acquisition, T.I. and J.U.

## DATA AVAILABILITY

The raw data supporting the conclusions of this article will be made available by the authors, without undue reservation.

## REFERENCES

1. Düzel, E., Penny, W.D., and Burgess, N. (2010). Brain oscillations and memory. Curr. Opin. Neurobiol. 20, 143–149.

2. Hsieh, L.T., and Ranganath, C. (2014). Frontal midline theta oscillations during working memory maintenance and episodic encoding and retrieval. Neuroimage 85, 721–729. Available at: 10.1016/j.neuroimage.2013.08.003.

3. Sestieri, C., Corbetta, M., Romani, G.L., and Shulman, G.L. (2011). Episodic memory retrieval, parietal cortex, and the default mode network: Functional and topographic analyses. J. Neurosci. 31, 4407–4420.

4. Menon, V. (2023). 20 years of the default mode network: A review and synthesis. Neuron 111, 2469–2487. Available at: 10.1016/j.neuron.2023.04.023.

5. Kaplan, R., Bush, D., Bonnefond, M., Bandettini, P.A., Barnes, G.R., Doeller, C.F., and Burgess, N. (2014). Medial prefrontal theta phase coupling during spatial memory retrieval. Hippocampus 24, 656–665.

6. Kerrén, C., Linde-Domingo, J., Hanslmayr, S., and Wimber, M. (2018). An Optimal Oscillatory Phase for Pattern Reactivation during Memory Retrieval. Curr. Biol. 28, 3383–3392.e6.

7. Ozawa, M., Davis, P., Ni, J., Maguire, J., Papouin, T., and Reijmers, L. (2020). Experience-dependent resonance in amygdalo-cortical circuits supports fear memory retrieval following extinction. Nat. Commun. 11, 1–16. Available at: 10.1038/s41467-020-18199-w.

8. Hebscher, M., Meltzer, J.A., and Gilboa, A. (2019). A causal role for the precuneus in network-wide theta and gamma oscillatory activity during complex memory retrieval. Elife 8, 1–20.

9. Watrous, A.J., Tandon, N., Conner, C.R., Pieters, T., and Ekstrom, A.D. (2013). Frequency-specific network connectivity increases underlie accurate spatiotemporal memory retrieval. Nat. Neurosci. 16, 349–356.

10. Ideriha, T., and Ushiyama, J. (2024). Behavioral fluctuation reflecting theta LJ rhythmic activation of sequential working memory. Sci. Rep., 1–11. Available at: 10.1038/s41598-023-51128-7.

11. VanRullen, R. (2018). Attention Cycles. Neuron 99, 632–634.

12. Helfrich, R.F., Fiebelkorn, I.C., Szczepanski, S.M., Lin, J.J., Parvizi, J., Knight, R.T., and Kastner, S. (2018). Neural Mechanisms of Sustained Attention Are Rhythmic. Neuron 99, 854–865.e5.

13. Landau, A.N., and Fries, P. (2012). Attention samples stimuli rhythmically. Curr. Biol. 22, 1000– 1004.

14. ter Wal, M., Linde-Domingo, J., Lifanov, J., Roux, F., Kolibius, L.D., Gollwitzer, S., Lang, J., Hamer, H., Rollings, D., Sawlani, V., et al. (2021). Theta rhythmicity governs human behavior and hippocampal signals during memory-dependent tasks. Nat. Commun. 12, 1–15.

15. Niv, Y. (2021). The primacy of behavioral research for understanding the brain. Behav. Neurosci. 135, 601–609.

16. Cowan, N. (2001). The magical number 4 in short-term memory: A reconsideration of mental storage capacity. Behav. Brain Sci. 24, 87–114.

17. Brainard, D.H. (1997). The Psychophysics Toolbox. Spat. Vis. 10, 433–436.

18. Dawson, M.R.W. (1988). Fitting the ex-Gaussian equation to reaction time distributions. Behav. Res. Methods, Instruments, Comput. 20, 54–57.

19. Hockley, W.E. (1984). Analysis of response time distributions in the study of cognitive processes. J. Exp. Psychol. Learn. Mem. Cogn. 10, 598–615.

20. Ratcliff, R., and Murdock, B.B. (1976). Retrieval processes in recognition memory. Psychol. Rev. 83, 190–214.

21. Delorme, A., and Makeig, S. (2004). EEGLAB: An open source toolbox for analysis of single-trial EEG dynamics including independent component analysis. J. Neurosci. Methods 134, 9–21.

22. Wobbrock, J.O., Findlater, L., Gergle, D., and Higgins, J.J. (2011). The Aligned Rank Transform for nonparametric factorial analyses using only ANOVA procedures. Conf. Hum. Factors Comput. Syst. - Proc., 143–146.

23. Shaffer, J.P. (1986). Modified Sequentially Rejective Multiple Test Procedures. J. Am. Stat. Assoc. 81, 826–831. Available at: https://www.tandfonline.com/doi/abs/10.1080/01621459.1986.10478341.

24. Benjamini, Y., and Hochberg, Y. (1995). Controlling the false discovery rate□: A practical and powerful approach to multiple testing author (s): Yoav Benjamini and Yosef Hochberg Source□: Journal of the Royal Statistical Society. Series B (Methodological), Vol. 57, No. 1 (1995), Publi. J. R. Stat. Soc. 57, 289–300.

25. Berens, P. (2009). CircStat: A MATLAB Toolbox for Circular Statistics. J. Stat. Softw. 31, 1–21.

26. Polich, J. (2007). Updating P300: An integrative theory of P3a and P3b. Clin. Neurophysiol. 118, 2128–2148.

27. Hillyard, S.A., Vogel, E.K., and Luck, S.J. (1998). Sensory gain control (amplification) as a mechanism of selective attention: Electrophysiological and neuroimaging evidence. Philos. Trans. R. Soc. B Biol. Sci. 353, 1257–1270.

28. Canavier, C.C. (2015). Phase-resetting as a tool of information transmission. Curr. Opin. Neurobiol. 31, 206–213. Available at: 10.1016/j.conb.2014.12.003.

29. Fiebelkorn, I.C., Saalmann, Y.B., and Kastner, S. (2013). Rhythmic sampling within and between objects despite sustained attention at a cued location. Curr. Biol. 23, 2553–2558. Available at: 10.1016/j.cub.2013.10.063.

30. Abdalaziz, M., Redding, Z. V., and Fiebelkorn, I.C. (2023). Rhythmic temporal coordination of neural activity prevents representational conflict during working memory. Curr. Biol. 33, 1855–1863.e3. Available at: https://www.biorxiv.org/content/10.1101/2022.12.02.518876v1 https://www.biorxiv.org/content/10.1101/2022.12.02.518876v1.abstract.

31. Pomper, U., and Ansorge, U. (2021). Theta-Rhythmic Oscillation of Working Memory Performance. Psychol. Sci. 32, 1801–1810.

32. Chota, S., Leto, C., van Zantwijk, L., and Van der Stigchel, S. (2022). Attention rhythmically samples multi-feature objects in working memory. Sci. Rep. 12, 14703. Available at: https://www.nature.com/articles/s41598-022-18819-z.

33. Peters, B., Kaiser, J., Rahm, B., and Bledowski, C. (2021). Object-based attention prioritizes working memory contents at a theta rhythm. J. Exp. Psychol. Gen. 150, 1250–1256. Available at: http://doi.apa.org/getdoi.cfm?doi=10.1037/xge0000994.

34. Cowan, N. (2010). The magical mystery four: How is working memory capacity limited, and why? Curr. Dir. Psychol. Sci. 19, 51–57.

35. Luck, S.J., and Vogel, E.K. (1997). The capacity of visual working memory for features and conjunctions. Nature 390, 279–281. Available at: https://www.nature.com/articles/36846.

36. Storm, R.W., and Pylyshyn, Z.W. (1988). Tracking multiple independent targets: Evidence for a parallel tracking mechanism. Spat. Vis. 3, 179–197.

37. Kaufman, E.L., Lord, M.W., Reese, T.W., and Volkmann, J. (1949). The Discrimination of Visual Number. Am. J. Psychol. 62, 498. Available at: https://www.jstor.org/stable/1418556?origin=crossref.

38. Kutter, E.F., Dehnen, G., Borger, V., Surges, R., Mormann, F., and Nieder, A. (2023). Distinct neuronal representation of small and large numbers in the human medial temporal lobe. Nat. Hum. Behav. 7, 1998–2007.

39. Baddeley, A.D., and Hitch, G. (1974). Working memory. Psychol. Learn. Motiv. - Adv. Res. Theory 8, 47–89.

40. Martín-Buro, M.C., Wimber, M., Henson, R.N., and Staresina, B.P. (2020). Alpha Rhythms Reveal When and Where Item and Associative Memories Are Retrieved. J. Neurosci. 40, 2510– 2518. Available at: https://www.jneurosci.org/lookup/doi/10.1523/JNEUROSCI.1982-19.2020.

41. Hayama, H.R., Vilberg, K.L., and Rugg, M.D. (2012). Overlap between the neural correlates of cued recall and source memory: Evidence for a generic recollection network? J. Cogn. Neurosci. 24, 1127–1137.

42. Rugg, M.D., and Vilberg, K.L. (2013). Brain networks underlying episodic memory retrieval. Curr. Opin. Neurobiol. 23, 255–260. Available at: 10.1016/j.conb.2012.11.005.

43. Varela, F., Lachaux, J.P., Rodriguez, E., and Martinerie, J. (2001). The brainweb: Phase synchronization and large-scale integration. Nat. Rev. Neurosci. 2, 229–239.

44. Fell, J., and Axmacher, N. (2011). The role of phase synchronization in memory processes. Nat. Rev. Neurosci. 12, 105–118.

45. Melby-Lervåg, M., and Hulme, C. (2013). Is working memory training effective? A meta-analytic review. Dev. Psychol. 49, 270–291.

